# Direct visualization of lysosome-mediated intracellular transport and membrane anchoring of TNF-α in human T cells

**DOI:** 10.64898/2026.10.07.757379

**Authors:** Alyssa Burgess, Hirushi Gunasekara, James Xie, Ying S. Hu

## Abstract

Tumor necrosis factor-alpha (TNF-α) represents a major pro-inflammatory cytokine associated with chronic inflammation. Enhanced understanding of intracellular trafficking of TNF-α enables the identification of drug targets that modulate TNF-α secretion at the cellular source. However, conventional cytokine assays provide only bulk measurements of secreted TNF-α or static measurements in fixed cells and therefore cannot provide mechanistic information. Here, we report a fluorescence imaging platform to visualize and quantify intracellular TNF-α trafficking in human T cells. Using a Jurkat T cell line stably expressing TNF-α fused to mStayGold as a model system, we revealed through single-particle tracking (SPT) that T cell activation increased the motility of TNF-α vesicles and that microtubule perturbation disrupted this increased motility. 3D structured illumination microscopy uncovered spatial associations between TNF-α vesicles and lysosomes, and SPT further demonstrated their coordinated transport. Using total internal reflection fluorescence microscopy, we captured the sub-second splitting of single TNF-α vesicles into multiple puncta that subsequently separated, suggesting delivery of TNF-α from the vesicle to the plasma membrane. Confocal-based super-resolution microscopy further resolved TNF-α vesicles fusing with the plasma membrane and transmembrane TNF-α. Together, these imaging capabilities for tracking and super-resolving the intracellular transport and membrane delivery of TNF-α in live T cells enable mechanistic studies of TNF-α trafficking and release.

## Introduction

TNF-α is a central mediator of inflammation and has been implicated in a wide range of inflammatory disorders, such as rheumatoid arthritis, Crohn’s disease, psoriasis, and inflammatory bowel disease (*1*, *2*). Targeting TNF-α has achieved clinical success in managing symptoms associated with chronic inflammation (*3*, *4*). Current therapeutic strategies primarily rely on neutralizing TNF-α with monoclonal antibodies, decoy receptor constructs that block TNF-α binding to its receptor, and Janus kinase inhibitors that block downstream TNF-α signaling. A major limitation of these approaches is the resulting broad immunosuppression, which disrupts the immune regulatory function of TNF-α. An alternative strategy is to block pathogenic TNF-α at its cellular source before secretion. However, the intracellular trafficking steps preceding TNF-α secretion remain poorly understood. Current cytokine quantification techniques provide only limited access to these intracellular processes. ELISA measures secreted TNF-α in the absence of the cellular context, whereas flow cytometry quantifies intracellular cytokine levels with limited or no spatial or temporal resolution. This limitation represents a major bottleneck for the development of cell-targeted approaches in battling chronic inflammation.

TNF-α secretion requires tightly regulated intracellular transport from sites of synthesis, packaging, and delivery to the plasma membrane, a process that likely involves membrane-enclosed vesicles (*5*). Previous work in macrophages has shown that TNF-α exits the trans-Golgi network through specialized tubular and vesicular carriers and is subsequently transported to the cell membrane, including the phagocytic cup (*6–8*). These studies suggest that lipopolysaccharide-induced activation can enhance TNF-α trafficking through post-Golgi compartments and promote its delivery to the cell surface in innate immune cells (*6*, *7*). Moreover, TNF-α is synthesized as the membrane-enclosed precursor pro-TNF-α, which is subsequently transported to the membrane, then cleaved by TNF-α-converting enzyme (TACE) to release soluble TNF-α (*9–11*). Although the molecular pathways regulating TNF-α production and extracellular signaling have been extensively characterized in innate immune cells, less is known about the mechanisms governing TNF-α vesicle trafficking in T cells. Studies in activated T cells have shown that TNF-α is transported through a multidirectional secretory pathway distinct from the polarized secretion of other cytokines (*12*). More recently, TNF-α vesicles in primary human CD4^+^ T cells have been demonstrated to interact with lysosomes and late endosomes, with these interorganelle interactions contributing to the transport of cytokine vesicles (*13*). While these studies establish that TNF-α trafficking in T cells involves interactions with the endosomal network, the dynamics of individual TNF-α vesicles, their interorganelle interactions in response to T cell activation, and their interaction with the plasma membrane for pro-TNF-α, remain poorly characterized.

To this end, fluorescence microscopy offers both spatial and temporal resolution, but diffraction-limited imaging in fixed cells using immunofluorescence staining cannot capture intracellular localization and dynamic transport (*14–16*). High spatial and temporal resolution enables the delineation of TNF-α subpopulations and their interorganelle dynamics responsible for transport. TNF-α exists as both vesicle-associated intracellular cargo (*6*, *13*) and transmembrane TNF-α (*17*, *18*), occupying distinct subcellular regions. In this regard, single-particle tracking (SPT) based on highly inclined and laminated optical sheet (HILO) illumination enables characterization of individual intracellular TNF-α vesicles, as it has been used to study endosomal trafficking, viral transport, and organelle movement (*19–24*). To investigate the volumetric distribution of TNF-α and its association with intracellular transport components, 3D structured illumination microscopy (SIM) represents an effective approach, as we previously demonstrated for interleukin-2 (IL-2) (*22*). For capturing membrane anchoring of TNF-α from its transport vesicles, total internal reflection fluorescence (TIRF) microscopy offers increased temporal resolution and high signal-to-noise imaging at a defined illumination and focal plane. For volumetric characterization of membrane loading of TNF-α, a confocal-based super-resolution technique with ultrahigh sensitivity, low photobleaching, and *z-*sectioning capability is required.

Here, we integrate these imaging modalities to characterize TNF-α trafficking in T cells. Using a stable Jurkat T-cell reporter expressing TNF-α fused to mStayGold as a model system, we characterize individual TNF-α vesicle motility before and after T cell activation and their vesicle trajectories, resolve their transport mechanism in relation to lysosomes and microtubules, and capture vesicular fusion with the plasma membrane to deliver transmembrane TNF-α. Together, this integrated imaging platform bridges dynamic tracking and 3D structural information to reveal TNF-α transport and release mechanisms.

## Results

### SPT reveals T cell activation upregulates transport dynamics of TNF-α

To investigate TNF-α transport dynamics in T cells, we used Jurkat T cells (E6.1) as a model system. We generated a Jurkat T-cell line stably expressing mStayGold fused to the c-terminus of TNF-α (TNF-α-mSG Jurkat T cells; Methods). TNF-α-mSG exhibited a predominantly punctate intracellular distribution, and we validated the phenotype using immunofluorescence (IF) staining. The white arrowheads in the boxed region in Fig. 1A outline the TNF-α puncta that overlapped between the mSG and IF channel. Minor differences between the two channels are likely attributed to endogenous TNF-α and antibody accessibility to intravesicular TNF-α during IF staining. Moreover, TNF-α-mSG puncta colocalized with intracellular DiD staining (Fig. 1B-C), supporting the notion that TNF-α-mSG is within membrane-enclosed vesicles.

**Figure 1.**
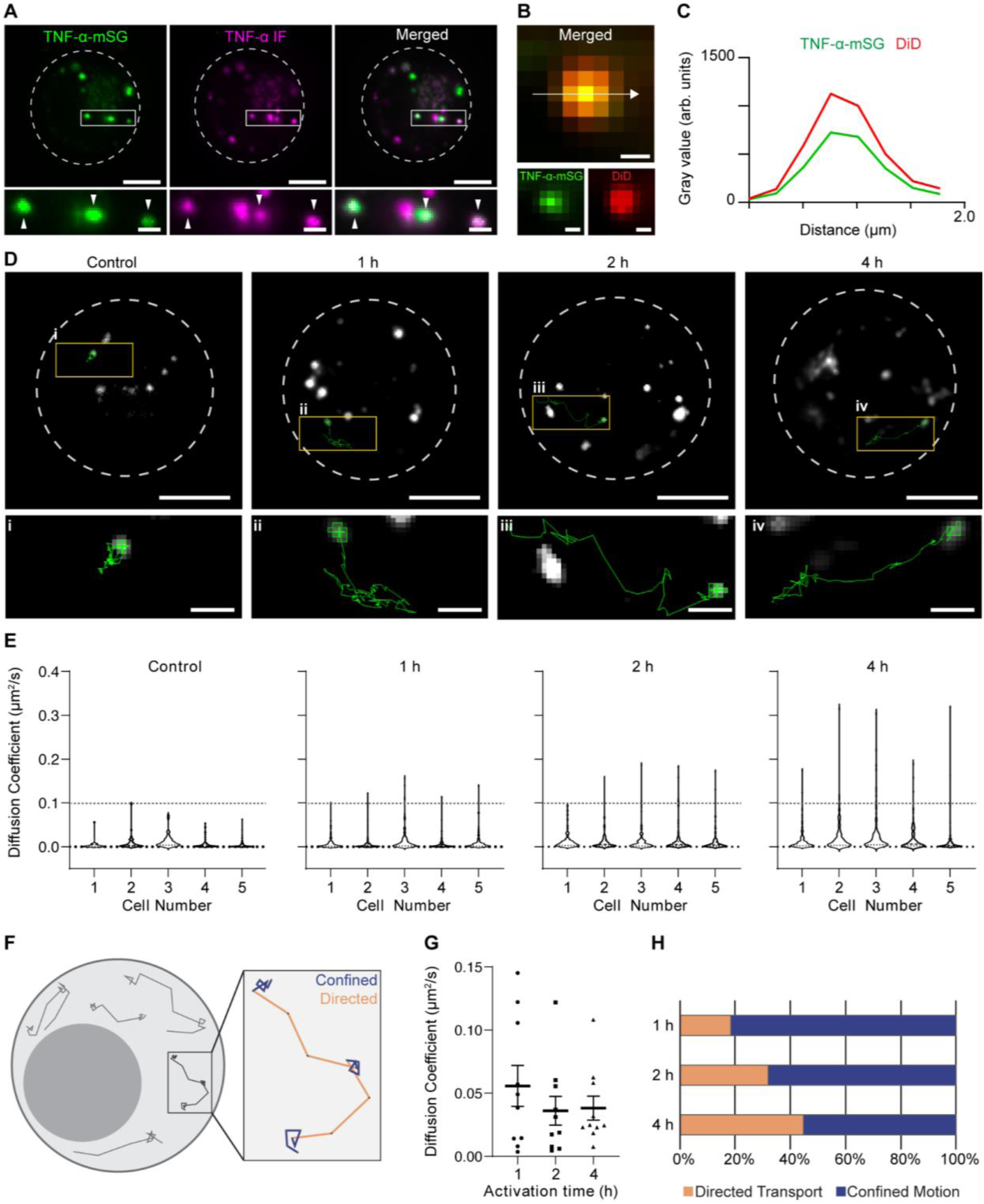
Single-particle tracking reveals increased directed transport of TNF-α vesicles after T cell activation. **(A)** Dual-color SIM (CrestOptics) images of unstimulated (control) TNF-α-mSG Jurkat T cells (left, green) immunostained for endogenous TNF-α (middle, magenta), and the merged image (right). Colocalizations appear in white. Arrowheads indicate colocalized TNF-α-mSG positive puncta with TNF-α immunostaining. **(B)** Representative single TNF-α-mSG puncta (green) colocalizing with DiD labeling of intracellular vesicles (red). Colocalization appears in yellow. **(C)** Cross-sectional line profile across the arrow shown in panel (B) illustrating the overlapping fluorescent signals. TNF-α-mSG (solid green line) and DiD (solid red line). **(D)** Representative single-particle tracks of TNF-α vesicles in control and Jurkat T cells activated for 1, 2, or 4 h with ImmunoCult. One selected TNF-α-mSG vesicle trajectory traced in green is shown at the top, and the magnified views of the trajectories (i-iv) are shown at the bottom. **(E)** Comparison of the diffusion coefficients of individual TNF-α-mSG vesicles from control T cells activated for 1, 2, or 4 h with ImmunoCult. *n=5* cells from 2 independent experiments.

We next investigated how T-cell activation influences TNF-α vesicle trafficking. TNF-α-mSG Jurkat T cells were activated with ImmunoCult Human CD3/CD28 T Cell Activator for 1, 2, or 4 h, and the vesicular dynamics were captured using HILO illumination. SPT was used to resolve individual TNF-α-mSG vesicle trajectories (Fig. 1D), and we calculated diffusion coefficients (*D*) from TNF-α-mSG vesicle trajectories for individual cells (Methods; Fig. 1E). Representative trajectories revealed an increase in TNF-α vesicle motilities after stimulation (Fig. 1D). TNF-α vesicles in unstimulated (control) cells exhibited predominantly confined trajectories, whereas activation resulted in more directional transport trajectories. In control Jurkat T cells, diffusion coefficients were 7.08 ×10^-3^ ± 3.54 ×10^-3^ μm²/s (mean ± SD), with maximum values approaching 0.1 μm²/s (Fig. 1E). While 1 h of activation resulted in more trajectories in directional transport (Fig. 1D), the mean diffusion coefficients remained relatively unchanged: 8.73 ×10^-3^ ± 3.40 ×10^-3^ (mean ± SD) for 1 h and 7.08 ×10^-3^ ± 3.54 ×10^-3^ μm²/s (mean ± SD) for control cells. However, following 2 and 4 h activation, the diffusion coefficients increased to 1.28 ×10^-2^ ± 1.25 ×10^-3^ (mean ± SD) and 1.73 ×10^-2^ ± 6.19 ×10^-3^ μm²/s (mean ± SD), respectively. These results demonstrate that T-cell receptor engagement upregulates TNF-α trafficking in T cells.

Inspection of individual trajectories revealed that TNF-α vesicles transitioned between both confined and directed transport states (Fig. 1F). The directed transport state likely reflects engagement of TNF-α vesicles with cytoskeletal transport machinery, whereas the confined state may represent periods of cargo sorting, pausing, or directional switching. To investigate whether the increase in the overall TNF-α vesicle dynamics resulted from faster movement during directed transport, *i.e.,* speed of the motor protein, or from overall increased engagement in the directed transport state compared to the confined state, we isolated and analyzed the directed transport segments from the entire tracks at 1, 2, and 4 h activation. Fig. 1G shows that the diffusion coefficient of directed transport segments remained relatively constant, with a wider distribution at the early time points. This observation suggests that T-cell activation did not alter the intrinsic mobility of TNF-α vesicles engaged in directed transport. However, Fig. 1H shows that TNF-α vesicles spent a greater portion of the track time in directed transport as activation duration increased. These results suggest that T-cell activation enhances TNF-α vesicle trafficking primarily by increasing the vesicle engagement in directed transport, positioning TNF-α for long-range intracellular trafficking.

The dashed line at 0.1 μm²/s indicates the maximum diffusion coefficient for control cells. **(F)** Schematic illustration of a single-particle track consisting of directed (orange) and confined (blue) transport segments. **(G)** Comparison of diffusion coefficients calculated from directed transport segments from activated T cells. Bars represent the mean and the standard error (SEM). *n=10* trajectories from 3 cells and 2 independent experiments. **(H)** Comparison of the fraction of total trajectory time spent in directed transport states across activation conditions. Horizontal bars represent the proportion of trajectory time classified as directed transport with respect to confined motion. Scale bars: 5 μm (A, D top), 0.5 μm (B), 1 μm (D bottom).

### TNF-α vesicles associate with the microtubule network for trafficking

The sustained, directed trajectories of TNF-α vesicles frequently exhibited a stop-and-go pattern, suggesting that their long-range movement may involve cytoskeleton-associated transport machinery (*25*, *26*). Given that microtubules serve as major tracks for long-range intracellular cargo transport (*27*, *28*), we next investigated the spatial relationship between TNF-α vesicles and the microtubule network in T cells. We immunostained α-tubulin in TNF-α-mSG Jurkat T cells. Fig. 2A shows the spatial association between TNF-α vesicles and microtubules. Interestingly, the spatial association took the form of adjacency rather than direct overlap.

**Figure 2.**
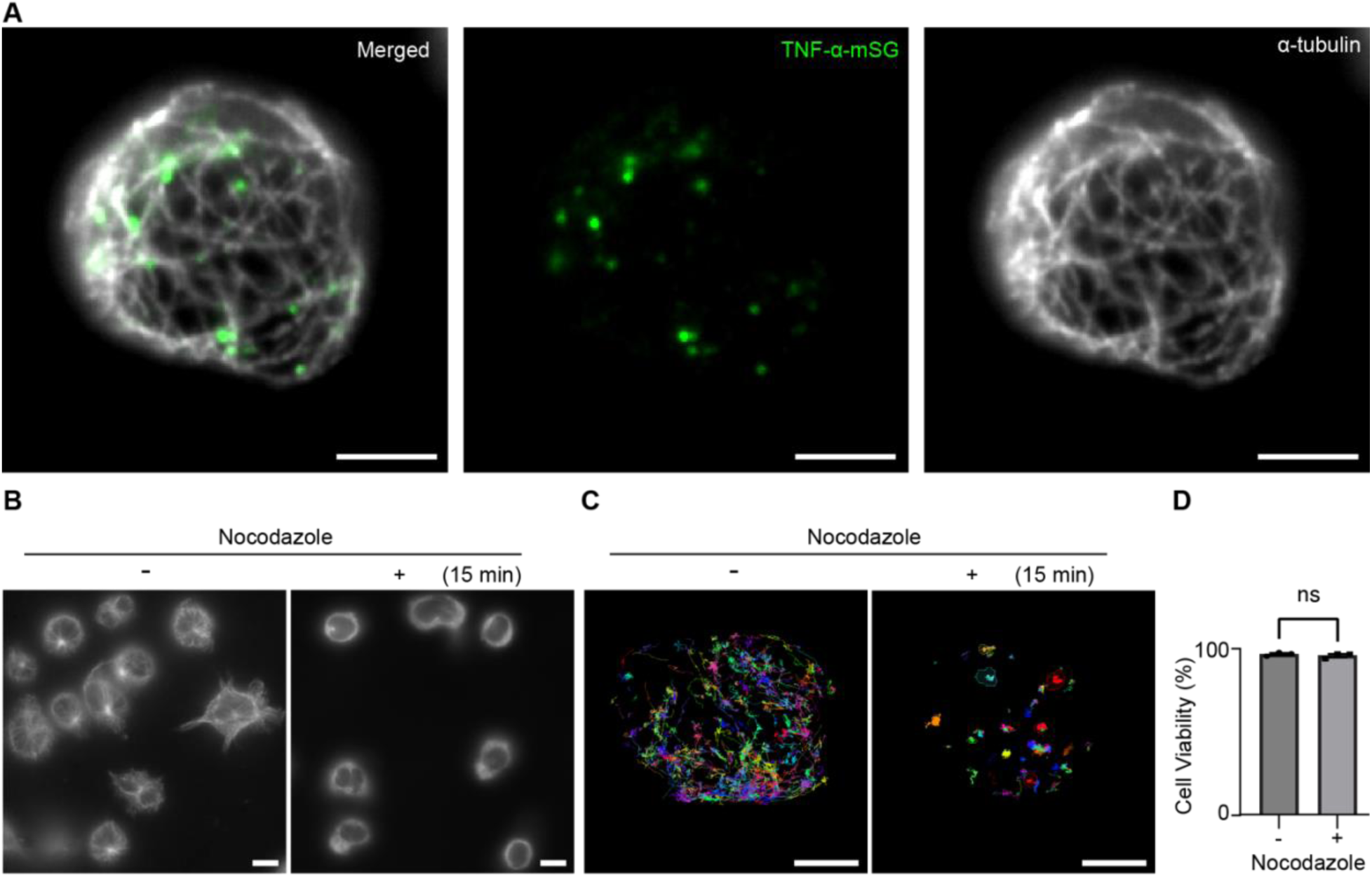
TNF-α vesicles associate with the microtubule network, and disruption of this network abrogates directed transport of TNF-α. **(A)** Representative merged image of a TNF-α-mSG Jurkat T cell immunostained for α-tubulin. **(B)** Representative images of α-tubulin in untreated and nocodazole-treated (15 min) Jurkat T cells. Cells were activated for 2 h using ImmunoCult. **(C)** Representative TNF-α-mSG single-particle tracks from untreated and treated Jurkat T cells activated by ImmunoCult for 2h. Each trajectory is distinguished by a unique color. **(D)** Comparison of the cell viability following nocodazole treatment assessed by Trypan blue exclusion. (*n=3* biological replicates; *p*>0.05). Scale bars: 5 μm (A, C) and 10 μm (B).

To assess the contribution of microtubules to TNF-α vesicle transport, we disrupted microtubule organization in activated TNF-α-mSG Jurkat T cells using nocodazole, a microtubule-depolymerizing agent. Epifluorescence imaging confirmed a substantial disruption of the microtubule network after 15 min of nocodazole treatment (Fig. 2B). SPT analysis subsequently revealed the loss of directed transport of TNF-α-mSG vesicle trajectories following nocodazole treatment (Fig. 2C). Cell viability remained relatively unchanged in both untreated and nocodazole-treated cells across independent experiments (Fig. 2D). Our results demonstrate that an intact microtubule network is required for the directed transport observed in activated T cells and support a microtubule-dependent mechanism for the long-range intracellular transport of TNF-α vesicles.

### TNF-α vesicles travel in tandem with late endosomal/lysosomal vesicles

T-cell cytokines have been previously shown to associate with components of the endosomal trafficking network during intracellular transport (*12*, *13*). In particular, TNF-α vesicles have been reported to associate with LAMP1-positive late endosomal/lysosomal vesicles (*13*). To investigate whether T cells utilize lysosomes as a mechanism for TNF-α transport, we first performed dual-color SPT of TNF-α-mSG vesicles and lysosomes labeled with LysoTracker Deep Red. In Jurkat T cells activated with ImmunoCult for 2 h, time-lapse imaging revealed coordinated movement between TNF-α vesicles and lysosomes (Fig. 3A, B). Time-lapse imaging frames further demonstrate that TNF-α vesicles remain juxtaposed to lysosomes throughout their movements. These observations suggest that TNF-α vesicles engage and co-traffic with lysosomes rather than transiently encountering them. The observation is also consistent with the involvement of microtubule networks in trafficking mechanisms, as lysosomes are known to be transported along microtubules (*26*, *29*).

**Figure 3.**
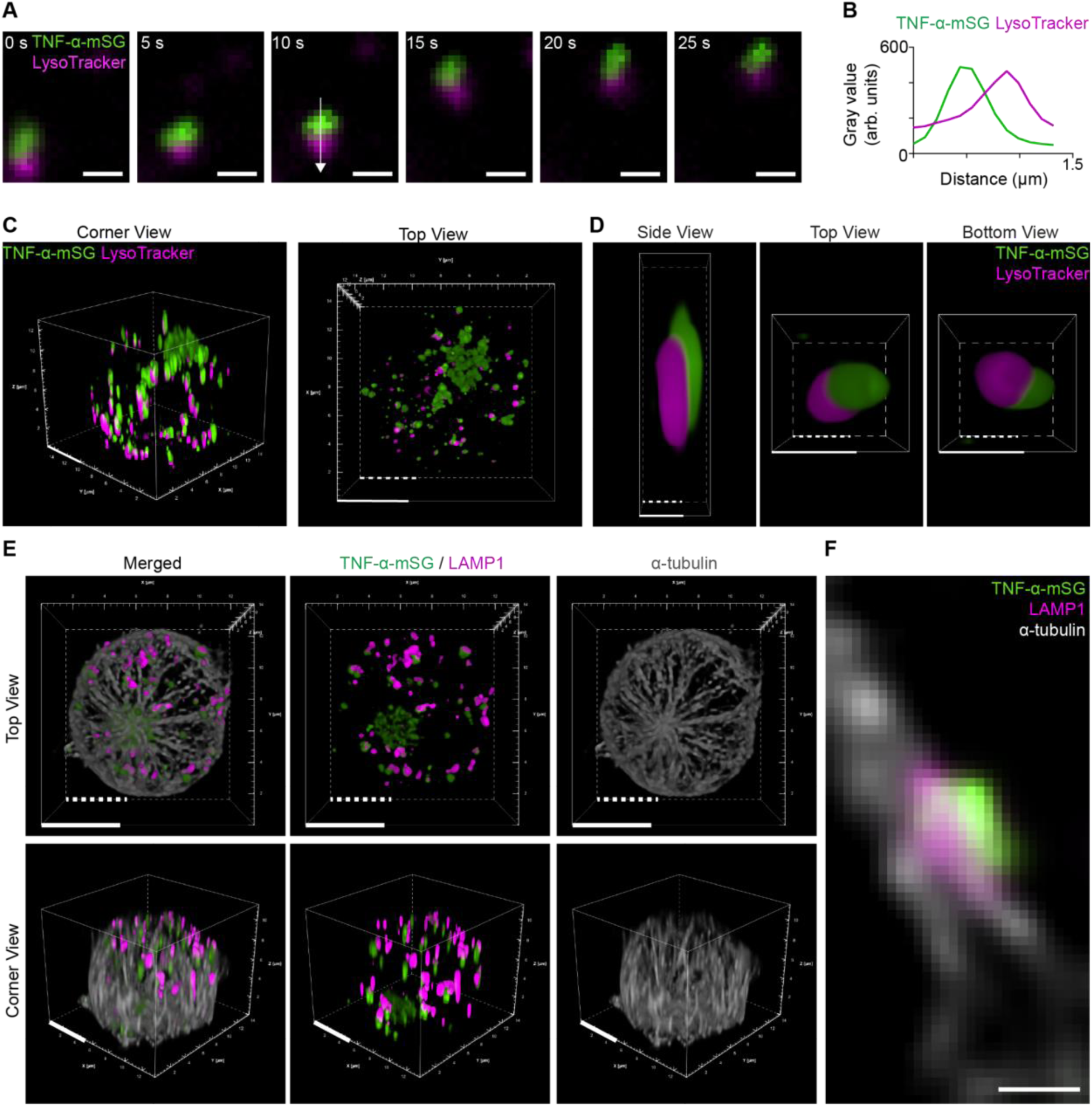
TNF-α vesicles hitchhike on lysosomes for coordinated transport along microtubules. **(A)** Representative images from dual-color SPT showing coordinated movement of TNF α-mSG vesicles (green) with lysosomal vesicles labeled with LysoTracker Deep Red (magenta). **(B)** Cross-sectional line profile across the indicated line in panel A depicting relative fluorescent intensity across the TNF-α-mSG vesicle and lysosome, showing juxtaposition of the signals instead of colocalization. TNF-α-mSG (green solid line) and lysosome (magenta solid line). **(C)** A representative 3D SIM image of TNF-α-mSG Jurkat T cells labeled with LysoTracker Deep Red. TNF-α-mSG and LysoTracker Deep Red are shown in green and magenta, respectively. An orthogonal view, ‘corner view,’ is shown on the left, and the top-down view of the volumetric view is shown on the right. **(D)** Magnified orthographic views of a single TNF-α vesicle juxtaposed to a LysoTracker-laden vesicle. TNF-α-mSG and LysoTracker Deep Red are shown in green and magenta, respectively. **(E)** Representative 3D SIM images of a TNF-α-mSG Jurkat T cell, immunostained for lysosomes (with LAMP1) and microtubules (α-tubulin). All channel merged images are shown on the left, TNF-α-mSG and LysoTracker overlaid image is shown in the middle, and the α-tubulin channel is shown on the right. The top-down view of the volumetric rendering is shown on top, and an orthogonal view, ‘corner view,’ is shown on the bottom. **(F)** A representative magnified 2D cross-sectional slice from the 3D volumetric reconstruction, highlighting a microtubule in spatial proximity with an associated pair of a LAMP1-positive vesicle and a TNF-α-mSG vesicle. Scale bars: 0.5 μm (A, D, F) and 5 μm (C, E). For volumetric renderings, the solid and dashed white scale bars indicate the scale at the front and back planes of the 3D volume, respectively.

We further examined this spatial phenotype across the 3D cytosol using SIM. Orthographic views of whole-cell volume renderings of Jurkat T cells activated with ImmunoCult for 2 h revealed that TNF-α vesicles were in tandem with lysosomes throughout the cellular volume (Fig. 3C-D). Importantly, 3D SIM confirmed that TNF-α vesicles were spatially associated with, rather than enclosed within, lysosomal vesicles. We further super-resolved the spatial organization of TNF-α vesicles, LAMP1-positive late endosomes/lysosomes, and microtubules using 3D SIM (Fig. 3E-F). Fig. 3F reveals the colocalization between a LAMP1^+^ lysosome (magenta) and microtubules (grey), as well as a TNF-α vesicle (green) tethered to the lysosome (magenta). This organization confirms that the TNF-α vesicle is not fused with the lysosome but rather “hitchhikes” onto it for intracellular co-trafficking following T-cell activation.

### TNF-α vesicles rapidly fuse with the plasma membrane and deliver transmembrane TNF-α

Increased vesicle motility could potentially account for increased TNF-α secretion owing to the accelerated lysosome-mediated delivery. However, a limitation of the SPT and SIM data is that they do not differentiate the directionality of TNF-α trafficking (*i.e.*, anterograde versus retrograde). In addition, the kinetics of TNF-α transfer from the vesicle to the plasma membrane remain entirely unknown. To visualize membrane-anchoring events, we performed TIRF imaging of TNF-α-mSG Jurkat T cells activated with ImmunoCult for 2 h and subsequently deposited onto glass surfaces coated with anti-CD3/CD28 antibodies (1:1). Live-cell imaging captured large intracellular TNF-α-mSG puncta approaching the basal plasma membrane (Fig. 4A, 1s) and subsequently resolving into multiple smaller puncta (Fig. 4A, 2-8s). The resulting puncta maintain stable point-spread functions consistent with localization within the imaging plane, suggesting they were likely confined to a 2D surface against the glass. Subsequently, the fluorescence intensity associated with individual large puncta became redistributed among the multiple lower-intensity puncta (Fig. 4B), consistent with the dispersal of TNF-α-mSG within or along the plasma membrane following vesicle delivery. The summed fluorescence intensity of the original structure and resulting puncta only decreased moderately over time (Fig. 4B). Although photobleaching likely contributed to the modest decrease in total fluorescence intensity over the imaging period, the appearance and persistence of spatially distinct TNF-α-mSG puncta following these events support the redistribution of TNF-α-mSG across the plasma membrane. Under our imaging conditions, the fastest transition from a concentrated TNF-α-mSG punctum to dispersed membrane-associated structures occurred on a timescale of less than 1 s (Fig. 4C). These observations indicate that TNF-α redistribution at the plasma membrane can occur rapidly following vesicle arrival, providing a mechanism for the prompt surface presentation of transmembrane TNF-α. We further captured vesicle fusion with the plasma membrane from the cytosolic space using a Nikon AX confocal microscope equipped with a Nikon spatial array confocal (NSPARC) detector, which drastically reduces photobleaching and enhances resolution due to its ultra-low-noise detector array. To selectively label the plasma membrane and nearby vesicles, activated TNF-α-mSG T cells were briefly stained with DiD for 2 min. 3D NSPARC imaging revealed TNF-α-mSG vesicles closely apposed to the plasma membrane throughout the cell surface (Fig. 4D). The TNF-α vesicles apposed to the plasma membrane exhibited a DiD-stained membrane, similar to the observation made in Fig. 1B. The presence of these structures at multiple regions of the cell surface is consistent with multidirectional TNF-α trafficking and delivery (*12*).

**Figure 4.**
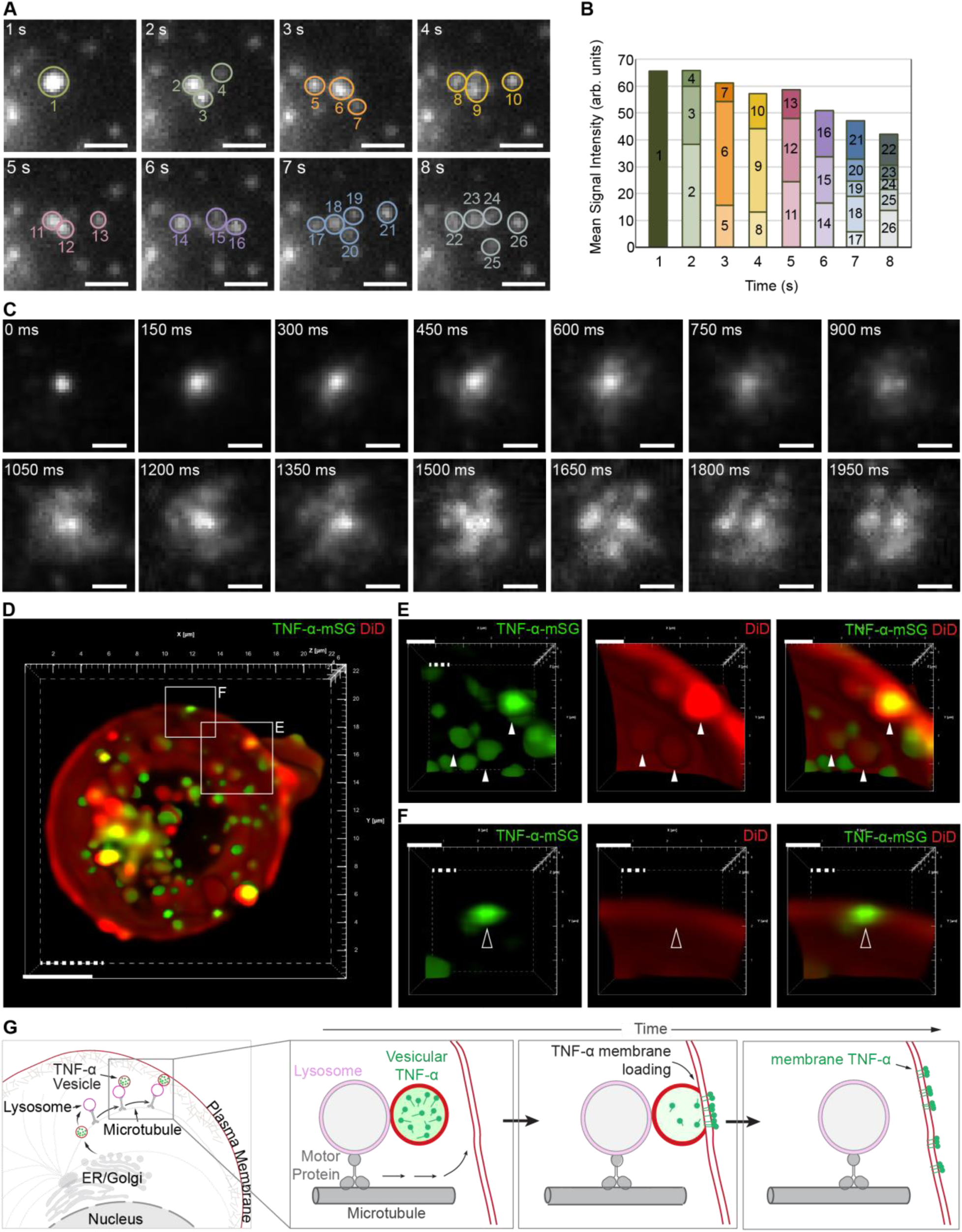
Plasma membrane anchoring of TNF-α captured by TIRF and confocal-based super-resolution imaging. **(A)** Time-lapse TIRF image sequence showing a single diffraction-limited TNF-α-mSG vesicle splitting into multiple smaller puncta that were spatially resolved as they diffused apart. Circles and the numbers indicate the initial TNF-α structure and resulting puncta used for fluorescence quantification. **(B)** Quantification of fluorescence intensity throughout the membrane-anchoring and splitting event. **(C)** Time-lapse TIRF image sequence showing a TNF-α-mSG vesicle undergoing sub-second splitting. **(D)** A representative cross-sectional view of a 3D NSPARC image of a TNF-α-mSG Jurkat T cell labeled with DiD. TNF-α-mSG and DiD are shown in green and red, respectively. **(E)** Magnified view of the boxed region indicated in panel D showing TNF-α-mSG vesicles enclosed in DiD-labeled vesicular membrane partially fused with the plasma membrane as indicated by solid white arrows. **(F)** Magnified view of the boxed region indicated in panel D showing a TNF-α-mSG signal directly associated with the plasma membrane without a discernible vesicular membrane, as indicated by the hollow arrowhead. **(G)** A schematic illustration of TNF-α vesicles approaching the plasma membrane, delivering the TNF-α onto the plasma membrane, and membrane-enclosed TNF-α subsequently diffuses apart on the plasma membrane. Scale bars: 1 μm (A, C, E, F) and 5 μm (D). For volumetric renderings, the solid and dashed white scale bars indicate the scale at the front and back planes of the 3D volume, respectively.

Intriguingly, the enhanced 3D spatial resolution and sensitivity enabled us to capture vesicles being fused with the plasma membrane, as shown by white arrowheads in Fig. 4E. We also observed TNF-α-mSG signal directly associated with the plasma membrane without a discernible surrounding vesicular membrane, consistent with incorporation of TNF-α into the plasma membrane following vesicle fusion (Fig. 4F, hollow arrows). Together with the rapid redistribution events captured by TIRF microscopy, these observations support a model in which membrane-enclosed TNF-α vesicles are transported to the cell periphery in coordination with lysosomes, engage the plasma membrane, and subsequently deliver TNF-α for incorporation and distribution within the plasma membrane (Fig. 4G). The membrane anchoring of TNF-α occurs rapidly, *i.e.*, within a second.

## Discussion

In this study, we integrated complementary imaging modalities to visualize and quantify TNF-α trafficking from the cytosol to the plasma membrane with high temporal and spatial resolution. SPT enables quantitative characterizations of individual TNF-α vesicle kinetics in response to T cell activation, while TIRF imaging captures rapid membrane anchoring of TNF-α. 3D super-resolution imaging with SIM and NSPARC provides a volumetric context for TNF-α vesicle interactions with lysosomes and the plasma membrane. Together, these approaches enable examining cytokine trafficking as a dynamic process.

We observed T-cell activation increases TNF-α vesicle dynamics primarily by increasing engagement in directed transport (Fig. 1F-H). Thus, activation appears to regulate how frequently or for how long TNF-α vesicles engage with the lysosomal transport machinery. Because the trajectory-derived diffusion coefficients do not distinguish anterograde from retrograde transport, future directional analyses using specific motor proteins will determine whether activation preferentially regulates movement toward or away from the plasma membrane. Our findings suggest that regulation of transport-state occupancy represents an additional layer through which T-cell activation can modulate cytokine trafficking. Rather than requiring changes to the existing transport machinery, increased cytokine dynamics can therefore arise from more sustained engagement with it. The dependence of directed transport trajectories on microtubule organization further supports this interpretation (Fig. 2). Microtubules provide major intracellular tracks for kinesin- and dynein-mediated long-range cargo transport (*25*, *33*). Nocodazole-mediated microtubule disruption substantially reduced long-range TNF-α trajectories and shifted vesicle behavior toward more confined motion, demonstrating that the directed transport identified by SPT requires an intact microtubule network.

Our studies further revealed a close spatial relationship between TNF-α vesicles and lysosomes. Live-cell imaging showed TNF-α vesicles maintaining close spatial association with LysoTracker-labeled vesicles (Fig. 3A-B), while 3D SIM resolved TNF-α vesicles adjacent to LAMP1-positive vesicles and microtubules (Fig. 3C). Importantly, TNF-α-mSG was not observed within the lysosomal lumen in our imaging. The observed trafficking behavior is consistent with our previous observations that T-cell activation increases the dynamics of acidic intracellular vesicles labeled with acridine orange and that IL-2 vesicles may be associated with LysoTracker-positive vesicles (*22*). This organization is also consistent with prior literature indicating that interorganelle tethering is a determinant of directional cytokine transport (*13*). Zhou *et al.* previously demonstrated using correlative light and electron microscopy that vesicles in human CD4^+^ T-cells remain physically associated with LAMP1-positive late endosomal/lysosomal vesicles while maintaining distinct membrane identities (*13*). Organelle “hitchhiking”, in which cargo associates with motile organelles to access their cytoskeletal transport machinery, has emerged as a mechanism for intracellular transport of cargoes, including RNA granules (*28*, *34*). Our observation of sustained TNF-α–lysosome association during bidirectional movement is consistent with this model, although the present imaging data do not establish that lysosomes directly drive TNF-α vesicle movement.

TIRF imaging captured rapid events in which larger TNF-α-mSG puncta approaching the basal membrane redistributed into multiple smaller membrane-associated structures (Fig. 4A, C). This redistribution was distinct from the previously described TNF-α release into the extracellular space, which was characterized by rather rapid dispersal of the fluorescence signal followed by its disappearance from the TIRF field (*13*). 3D NSPARC imaging resolved both membrane-enclosed TNF-α vesicles closely apposed to the plasma membrane (Fig. 4E) and TNF-α-mSG directly within the plasma membrane without a separately resolvable vesicular boundary (Fig. 4F). Together, these observations provide temporal and spatial evidence for sequential stages of coordinated TNF-α vesicle trafficking with lysosomes, and subsequent transport to the T cell membrane, its anchoring to the plasma membrane (Fig. 4D-G). SPT of TNF-α following membrane incorporation will also be required to establish whether it remains localized near the delivery site or subsequently redistributes across the plasma membrane. A limitation of our study is that TNF-α-mSG was ectopically expressed from a lentiviral construct under the control of a CMV promoter and therefore does not recapitulate endogenous TNF-α transcriptional regulation downstream of T-cell receptor activation. Overexpression may also alter TNF-α abundance, while fusion to mStayGold could influence cargo processing or transport. However, we expect the preservation of the TNF-α transport mechanism. In addition, while Jurkat E6.1 cells provide a useful model for quantitative live-cell imaging, they do not fully reproduce the physiology of primary T cells. As such, the present reporter system is best interpreted as a tool for resolving TNF-α-associated vesicle trafficking behaviors.

In summary, our study establishes an imaging platform that connects quantitative live-cell trajectory analysis with 3D structural information across multiple spatial and temporal scales. By combining TIRF microscopy, SPT, 3D SIM, and NSPARC imaging, TNF-α trafficking can be resolved from transitions between intracellular transport states and microtubule-dependent movement to organelle association and plasma membrane delivery. Beyond TNF-α, this imaging platform offers a strategy to investigate the intracellular transport of cytokines and other secretory cargoes whose dynamic behavior is difficult to capture with conventional endpoint measurements.

## Acknowledgements

The authors thank Dr. Yu-Shiuan Cheng and Prof. Alison Ondrus for assistance with NSPARC imaging. This work is supported by the National Institute of General Medical Sciences of the National Institutes of Health R35GM146786 (Y.S.H). The content is solely the responsibility of the authors and does not necessarily represent the official views of the National Institutes of Health.

## Author contributions

H.G., Y.S.H., and A.B. conceived the work, designed and planned the experiments. A.B. performed cell culture, lentiviral transduction, immunostaining, microscopy, data analysis, and visualization. J.X. performed MATLAB SPT analysis. A.B. and H.G. drafted the manuscript, and all reviewed and edited the manuscript.

## Data Availability Statement

The original data is available upon request to the corresponding author.

## Methods

### Materials and reagents

RPMI 1640 Medium (11875119), penicillin-streptomycin (15140–122), 10×DPBS (MT20030CV), LysoTracker Deep Red (L12492), TNF alpha Monoclonal Antibody (28401) (MA5-23720), alpha Tubulin Monoclonal Antibody (DM1A) (14-4502-82), Goat anti-Rabbit IgG (Heavy chain) Superclonal Secondary Antibody, Alexa Fluor 647 (A27040), Goat anti-Mouse Kappa Light chain Superclonal Secondary Antibody, Alexa Fluor 555 (A28180), Vybrant Multicolor Cell-Labeling Kit DiD solution (V22889), Trypan Blue (15250–061), bovine serum albumin (BSA) (BP1600-100) were purchased from Thermo Fisher Scientific. LAMP1 (D2D11) Rabbit Monoclonal Antibody (9091S) was purchased from Cell Signaling Technology. Jurkat E6.1 Cell line (88042803-1VL), FBS (F0926), Poly-L-lysine solution (P4707-50ML), Saponin from *Quillaja* sp. (S4521) were purchased from MilliporeSigma. 8-well chambered cover glasses (No. 1.5, C8-1.5H-N) were ordered from Cellvis. InVivoMAb anti-human CD3 (BE0001-2) and InVivoMAb anti-human/monkey CD28 (BE0291) were purchased from Bio X Cell. Paraformaldehyde (16% w/v aq. stock solution; 15710) was purchased from Electron Microscopy Sciences. ImmunoCult^TM^ Human CD3/CD28 T Cell Activator (10971) was purchased from STEMCELL Technologies.

### Cell culture

Jurkat E6.1 T cells and TNF-α-mSG Jurkat T cells were cultured in RPMI 1640 with 10% FBS and 100 U/mL penicillin-streptomycin. Cells were maintained at 37 °C in a humidified atmosphere of 5% CO_2_ and split upon reaching confluence.

### Generation of TNF-α-mSG reporter cell line

A lentiviral construct expressing human TNF-α fused to mStayGold at the C-terminus was obtained (VectorBuilder). Jurkat E6.1 T cells were transduced via spinoculation (800×g, 30 min) with 1 μg/mL polybrene. After 48 h, puromycin (0.5 μg/mL) was added to select for stable expression. Expression and proper localization of TNF-α-mSG were confirmed by immunofluorescence (Fig. 1A).

### T cell activation

TNF-α-mSG Jurkat T cells were counted, centrifuged at 300 ×*g* for 3 min, and then resuspended in fresh culture medium. A volume of 200 μL of T cells (∼100,000 cells) was added to a well of a 96-well plate. 2.5 μL of ImmunoCult was added to the cells in the 96-well plate. Cells were incubated for the indicated times (0-4 h) at 37°C with 5% CO_2_. After incubation, the cells were placed in 1.5 mL microcentrifuge tubes, centrifuged at 300 × g for 3 min, and then resuspended in fresh culture medium.

### Preparation of coated surfaces

An eight-well chambered coverglass was prepared with either poly-L-lysine (PLL) or anti-CD3/CD28. For PLL surfaces, 0.01% PLL solution was added to each well and incubated for 2 h at 37°C before use. For activating surfaces, wells were coated overnight at 4°C with anti-CD3 (OKT3, 5 μg/mL) plus anti-CD28 (5 μg/mL) in 1X PBS, prior to use.

### LysoTracker labeling

Activated TNF-α-mSG Jurkats (1×10^6^ cells/mL) were incubated with LysoTracker Deep Red (75 nM) in complete growth medium for 30 min at 37°C. Cells were washed by centrifugation (300×g, 3 min) and resuspended in prewarmed growth medium before landing and imaging. Labeled Jurkats were added to coated wells (50-80,000 cells/well) and allowed to land (15 min on PLL or 1 h on CD3/CD28 at 37°C).

### Immunofluorescence staining

Activated TNF-α-mSG cells were settled on coverslips, then fixed in 4% PFA for 15 min at room temperature and washed. Cells were subsequently permeabilized and blocked in 1X PBS with 0.05% saponin and 5% BSA for 1 h. The blocking buffer was replaced with primary antibodies against LAMP1 (1 μg/mL), α-tubulin (10 μg/mL), or TNF-α (25 μg/mL) and incubated overnight at 4°C. The secondary antibody (goat anti-mouse Alexa Fluor 555 (2 µg/mL) and/or goat anti-rabbit Alexa Fluor 647 (2 µg/mL) in blocking buffer) was added and incubated for 2 h in a dark, humid chamber. After incubation, the antibody solution was replaced with 1× DPBS supplemented with 0.05% saponin and was washed three times. Cells were maintained in 1× DPBS until imaging.

### Cell viability assay

Jurkat TNF-α-mSG cells were grown in complete RPMI media. Samples were either left untreated or treated with 20 mM Nocodazole and placed at 37°C for 15 min. After incubation, each solution was diluted with trypan blue to a final concentration of 1 mg/mL and incubated for 1 min at room temperature. 10 μL of this solution was loaded onto a hemocytometer and imaged immediately after. Percent viability was determined based on the number of identified live cells divided by the total number of cells counted.

### Microscopy

#### Epifluorescence Microscopy

Images were acquired using a Nikon TS-2R inverted microscope equipped with a 40x/0.65 objective (Plan OFN22 Ph2 DL) and a Moment camera (Serial Number: A23G635016). Images were collected with no binning, a 12-bit readout, and 50 ms exposure time.

#### TIRF Microscopy

TIRF imaging was performed using an inverted microscope (Nikon Instruments, Eclipse Ti2-E) with 100×/1.49 NA oil immersion objective (CFI Apochromat TIRF 100XC), equipped for TIRF imaging with a quad-band TIRF set TRF89901-ET-405/488/561/640 nm (Chroma Technology), and emission filters ET450/50 m, ET525/50 m, ET600/50 m, ET700/75 m (Chroma Technology). Samples were illuminated sequentially using 488 nm and 638 nm lasers at 10 and 2 W/cm^2^, respectively, with 50 ms integration times. Time-lapse imaging (5-15 min duration) with 150 ms or 1 s non-illuminating intervals (NII) was recorded using ND acquisition in Nikon NIS-Elements Advanced Research software (version 5.42.06) and a Prime 95B sCMOS camera (Photometrics, Serial Number: A21C203010) at 16-bit images, no pixel binning with an effective imaging pixel size of 110 nm. All live imaging was performed at room temperature.

#### 3D SIM Microscopy

SIM images were acquired using an inverted microscope (Nikon Instruments, Eclipse Ti2-E) equipped with the SIM system (DEEPSIM X-LIGHT, CrestOptics). A 60×/1.42 oil objective (Plan Apochromat λD 60OIL OFN 25 DIC N2, 108 nm image pixel size) was used to acquire images, with a Kinetix sCMOS camera (serial number: A23H723002). Image acquisitions were performed using NIS-Elements Advanced Research software (Version: 5.42.06). 3D SIM images of fixed Jurkat T cells were captured using the ‘Deep Imaging’ SIM mode with 488, 561, and 640 nm laser illumination, an integration time of 50 ms, and a step size of 0.5 μm. Captured z-stacks were reconstructed in NIS-Elements software, followed by blind 3D deconvolution-blending: depth-coded alpha. Image contrast of the 3D-rendered images was adjusted for each channel in the LUTs (Gamma value = 1.00).

#### 3D NSPARC Microscopy

NSPARC images were acquired using a Nikon AX Confocal Microscope System with an integrated NSPARC detector mounted on a Nikon Ti-2 Eclipse inverted microscope. A 60×/1.42 NA oil immersion objective was used with Galvano bidirectional scanning (dwell time of 0.5 µs) in NSPARC confocal (CF) mode at a resolution of 1,024 × 1,024 pixels, corresponding to an effective pixel size of 0.08 µm. Samples were illuminated with 488 nm and 638 nm lasers. Volumetric image stacks were acquired with a z-step size of 0.172 µm and were deconvolved using ‘3D Deconvolution’ in NIS-Elements Advanced Research software (version 6.20.02).

### Data processing and SPT

TIRF time-lapse movies and 3D SIM movies were created within NIS-Elements Advanced Research software (Version: 5.42.06), then exported. Representative tracking movies were created at 7 fps. Further visualization and analysis of TIRF microscopy images were performed using ImageJ (Fiji) 1.54 f.

#### Qualitative SPT for visualization

For visualization of the individual vesicular dynamics (Fig. 1C), 2D SPT was performed using the ‘Track Binaries’ analysis option in the 2D tracking module of NIS-Elements Advanced Research software (Version 5.21.03). TNF-α vesicles were selected based on the intensity, and binary masks were created (Lower intensity threshold value: 5000 and Maximum intensity threshold value: 65535). 2D objects were tracked using the ‘Random Motion’ model, at maximum speed, with intensity-based centroid calculation. A standard deviation multiplication factor of 3.5 was used to track individual objects. Tracks were linked with 5 maximum gaps allowed. Tracks with 5 or fewer frames were removed.

#### Quantitative SPT

Single particle trajectories were analyzed in MATLAB (R2022b). Raw TIRF movies were imported into MATLAB as .tiff files, and each frame was filtered for analysis. To remove background noise, each frame was processed with a bandpass filter with a particle diameter scale corresponding to 7 pixels or greater. Vesicles less than 7 pixels in diameter would be considered as background noise. Additionally, only particles above the threshold intensity parameter of 10 were considered. Note that vesicles with lower resolution or size require reducing the parameters. The filter, intensity threshold, and particle diameter enhanced the desired particle to generate clearer images that emphasize the lysosome and cytokine. Locations of the vesicles were identified from the filtered frames through peak detection utilizing the same parameters. The detected vesicles were subjected to subpixel localization through a 2D Gaussian fit. This allows for predicting the center of a vesicle with nanometer precision despite each pixel being significantly larger. This was computed for each detected vesicle in one frame, repeated for the entire file, and stored. To plot the points and trajectories, the stored localized coordinates underwent additional filtering to remove poor fits and nonphysical locations. Pixel sizes were calibrated to 110 nm per pixel to convert vesicle coordinates into spatial units. Undefined (NaN) and negative coordinates were excluded from the final data set. The spatial positions of each vesicle were plotted, and the trajectories were reconstructed using the nearest neighboring maximum displacement of 20 pixels. Note that if vesicles are greater than 20 pixels apart within consecutive frames, this would be considered as two separate trajectories. Utilizing the MATLAB brush tool, the trajectory of interest was selected and saved as a new MATLAB workspace variable for further analysis.

#### Statistical Analysis

Data were analyzed using GraphPad Prism (9.5.1). Comparisons between two groups were performed using an unpaired, two-tailed Student’s t test assuming equal variances. A *p* value < 0.05 was considered statistically significant.

#### Quantifying Mean square Displacement (MSD) and Diffusion coefficients (D)

The trajectory of interest was subjected to MSD and *D* computations based on Equation 1 and 2 for 2D SPT. MSD computations were performed for each time step of the entire trajectory, and the slope of the first 10 time steps were divided by 4 to quantify *D* in two dimensions.

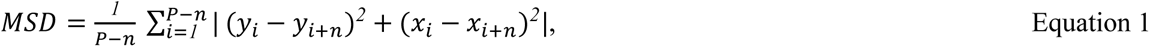

where *P* is the total number of points in a trajectory, and *n* is the lag time.

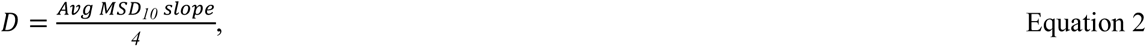

where *Avg MSD_10_ slope* is the slope for the first 10 points of the MSD plot, and the slope is calculated based on the MSD *vs.* lag time in frame.

#### Directed transport vs. full track Analysis

To characterize the directed transport of vesicles, segments of the trajectories were analyzed separately from their confined motion. Directed transport was identified as a period of sustained displacement occurring at any point along the entire trajectory. Only segments with 10 or greater consecutive points were considered. Each segment was required to begin at the final localized position of the initial region of confined motion or terminate at the first localized position at the final region of confined motion. Note that if a segment is between two regions of confined motion, both localized position conditions must be satisfied. For each segment of directional transport, MSD and *D* analysis were performed to characterize its biophysical dynamics.

